# Epigallocatechin Gallate (EGCG) as a Protective Agent Against Enzymatic Stromal Degradation in Caprine and Ovine Corneas: Towards Novel Therapeutics for Keratoconus

**DOI:** 10.64898/2026.01.30.697846

**Authors:** Muhammad Usman Ali Khan, Syed Tauheed Ahmad, Sabawoon Nisar, Robina Nematullah Khan, Faisal F. Khan

**Affiliations:** CECOS-RMI Precision Medicine Lab, RMI Laboratory Building, Peshawar, Pakistan; Trinity School, Lahore, Pakistan; Institute of Integrative Biosciences, CECOS University, Peshawar, Pakistan; Centre for Genomic Sciences, Rehman Medical Institute, Peshawar, Pakistan

## Abstract

**Purpose:** Keratoconus (KC) is a progressive corneal ectatic disorder marked by stromal thinning, enzymatic degradation, and oxidative stress. Conventional corneal collagen cross-linking (CXL) therapy poses risks. Epigallocatechin gallate (EGCG), a catechin found in green tea, has been shown to cross-link collagen, inhibit proteases, and modulate inflammation, suggesting its potential as an alternative therapy for KC. This study aimed to (1) evaluate the protective effects of EGCG against collagenase-mediated stromal degradation in caprine and ovine corneas and (2) investigate the use of caprine and ovine cornea as viable *ex vivo* models for corneal ectasia research.

**Methods:** Goat (n=4) and sheep (n=8) corneoscleral buttons were cultured in an air–liquid interface (ALI). Tissue viability was monitored by transparency grading and histology. The corneal ectasia environment was induced by collagenase type I digestion. An EGCG-rich extract was utilized for the treatments.

**Results:** Goat and sheep corneas remained intact for 7 days in MEM-based medium, though transparency decreased, with edema (whitening) by Day 7. Histology confirmed stromal loosening and reduced keratocyte cell density but preserved architecture sufficient for enzymatic digestion. EGCG pre-treatment and post-treatment exposure were associated with greater resistance to enzymatic digestion, with greater stromal preservation of quadrants in the EGCG pre-treated quadrant than in PBS pre-treated controls.

**Conclusions:** This study provides proof of concept that EGCG extract confers protection against enzymatic degradation in goat and sheep corneas, highlighting its potential as a protective agent for corneal ectatic disorders like Keratoconus. Additionally, goat and sheep corneas can represent practical and ethical *ex vivo* models for short-term ocular research. Future work should focus on cytotoxicity, biomechanical validation, optimized culture conditions, and *in vivo* studies.

## 1.0 Introduction

Keratoconus (KC) is a progressive corneal ectatic disorder characterized by thinning and protrusion of the central or paracentral cornea, leading to irregular astigmatism, myopia, and significant visual impairment, most often presenting in adolescence or early adulthood (Santodomingo-Rubido *et al*., 2022). Once considered a non-inflammatory degenerative disease, KC is now recognized as multifactorial in origin, with genetic and environmental contributions (Davidson *et al*., 2013; Singh *et al*., 2024). Epidemiological studies report a significant global variation in the prevalence and incidence rates of Keratoconus, with disproportionately higher rates in Middle Eastern and Asian populations (Santodomingo-Rubido *et al*., 2022).

The pathophysiology of KC involves both biochemical and structural dysregulation of the corneal stroma. Elevated levels of proteolytic enzymes, particularly collagenases and matrix metalloproteinases (MMPs), have been consistently detected in the tears and corneal tissues of KC patients, reflecting an imbalance between extracellular matrix degradation and protective inhibition. (Mackiewicz *et al*., 2006; Balasubramanian *et al*., 2012) Increased collagenase and gelatinase activity has been demonstrated in keratoconic corneal cultures, while levels of natural protease inhibitors are reduced, collectively accelerating stromal thinning and biomechanical weakening (Mackiewicz *et al*., 2006, Balasubramanian *et al*., 2012). In parallel, oxidative stress and elevated inflammatory mediators such as interleukin-1β (IL-1β), tumor necrosis factor-α (TNF-α), and interleukin-6 (IL-6) contribute to corneal degeneration, underscoring the dual biochemical and biomechanical drivers of disease progression (Lema *et al*., 2009; Balasubramanian *et al*., 2012; Sharif *et al*., 2018).

Currently, the only FDA-approved treatment shown to halt KC progression is corneal collagen cross-linking (CXL), which employs riboflavin and Ultraviolet A (UVA) light to induce additional covalent bonds within the stromal collagen (Wollensak *et al*., 2003; Papachristoforou *et al*., 2025). The standard epithelium-off Dresden protocol effectively stabilizes corneal biomechanics but is associated with complications such as postoperative pain, corneal haze, keratocyte loss, infectious keratitis, and endothelial damage. Transepithelial (“epi-on”) and accelerated modifications were developed to improve safety and patient comfort; however, these suffer from limited riboflavin penetration and reduced biomechanical efficacy. Complication rates include vision loss in ∼3% of treated eyes and failure rates of 7–8% (Papachristoforou *et al*., 2025; Shankar *et al*., 2024). Importantly, CXL cannot be performed on corneas thinner than 400 μm due to the risk of endothelial damage, excluding a significant proportion of advanced KC patients from treatment (Santodomingo-Rubido *et al*., 2022). Furthermore, while effective in halting ectatic progression, CXL does not address key disease mechanisms such as oxidative stress, inflammation, or ongoing enzymatic degradation (Mas Tur *et al*., 2017; Papachristoforou *et al*., 2025). These limitations highlight the need for novel therapeutic strategies with broader efficacy. Natural compounds such as epigallocatechin-3-gallate (EGCG), the major catechin in green tea (*Camellia sinensis*), represent promising candidates as alternative cross-linkers or adjunct therapies. Structurally, EGCG is a flavan-3-ol polyphenol with multiple hydroxyl groups and a galloyl moiety, conferring strong antioxidant activity and the ability to bind collagen fibrils via hydrogen bonding (Capasso *et al*., 2025). Importantly, EGCG has been directly shown to function as a collagen cross-linking agent in multiple non-corneal tissues. For example, in guided bone regeneration models, EGCG cross-linked collagen membranes exhibited increased cross-link density, fiber diameter, tensile strength, thermal stability, and resistance to enzymatic degradation, while preserving collagen backbone integrity (Chu *et al*., 2016). Similarly, in decellularized porcine osteochondral xenografts, EGCG treatment significantly enhanced collagen cross-linking, restored native mechanical properties, and conferred marked resistance to collagenase digestion (Elder *et al*., 2017).

Beyond structural reinforcement, EGCG inhibits collagenases and MMPs, reduces IL-1β-mediated pro-MMP-9 activation, and suppresses oxidative and inflammatory pathways implicated in KC (Cavet *et al*., 2011; Sapula *et al*., 2023). In ocular studies, EGCG has also promoted wound healing and reduced epithelial inflammation, further underscoring its therapeutic versatility (Urolita *et al*., 2024). These findings suggest that EGCG could simultaneously target multiple pathophysiological drivers of KC, indicating a holistic therapeutic profile.

To explore the potential role of EGCG in protecting against keratoconus-related stromal changes, appropriate experimental models are essential. Collagenase-mediated stromal digestion has been used extensively to simulate features of corneal ectasia, such as stromal thinning, steepening, and reduced biomechanical stability, in both human and animal corneas (Hong *et al*., 2012; Arafat *et al*., 2014). *Ex vivo* models provide a useful platform for corneal disease modelling by maintaining native stromal architecture and cell–matrix interactions while avoiding some of the ethical and logistical limitations of *in vivo* work. Goat and sheep corneas remain relatively underutilized in this context, despite their practical and biological advantages. Goat corneas are anatomically comparable to human corneas and can remain viable in culture for up to 15 days (Madhu *et al*., 2018). Sheep corneas include Bowman’s layer and display high protein sequence similarity to humans (≈90%) (Shankar *et al*., 2024). Their ready availability from abattoirs further supports their use as accessible *ex vivo* research tissues. In this study, we developed collagenase-induced stromal degradation models in goat and sheep corneas and carried out a preliminary assessment of EGCG extract treatment on stromal preservation.

## 2.0 Methods

### 2.1 Materials and Reagent Preparation

The transport medium consisted of phosphate-buffered saline (PBS, 1×, Invitrogen) supplemented with 1× penicillin–streptomycin and 1× amphotericin B (Gibco) to prevent bacterial and fungal contamination during eye collection and transport. For the culture media, Minimum Essential Medium (MEM, L-glutamine containing) (Gibco) was used as basal medium, supplemented with 10% fetal bovine serum (FBS) (Gibco), 1× penicillin– streptomycin, 1× amphotericin B (Gibco), and 10 ng/mL epidermal growth factor (EGF) (Gibco). MEM was used as the basal medium due to reagent availability constraints at the time of experimentation. Media were prepared fresh, filter-sterilized, and stored at 4 °C until use. Supporting domes for corneal stabilization were prepared separately from 1.5% agarose and 0.5% gelatin in PBS, autoclaved or sterile filtered before use. These domes provided a stable scaffold for maintaining corneal curvature during culture. These protocols were adapted from studies that cultured caprine, ovine, and equine corneas in an *ex vivo* setting. (Marlo *et al*., 2017; Madhu *et al*., 2018; Okurowska *et al*., 2024).

Collagenase solution was prepared from lyophilized collagenase type I (EMD Millipore) reconstituted in phosphate-buffered saline (PBS), with working concentrations freshly prepared immediately before each experiment. An EGCG-rich extract was prepared from Brazilian prime green tea using a two-step temperature-dependent solubilization protocol in double-distilled water as described by Bazinet *et al*. (2007), which selectively enriches catechins and yields higher EGCG content after the second extraction step. For corneal application, the concentrated extract was diluted in PBS (1×) as a physiologically compatible isotonic vehicle, filter-sterilized, and stored at 4 °C protected from light-induced degradation by wrapping in aluminum foil (Baek *et al*., 2021) until use. Based on the reported EGCG content of the extract (503.0 ± 47.2 µg/mL; Bazinet *et al*., 2007), this corresponds to an approximate EGCG concentration of ∼1.1 mM. A working solution corresponding to an estimated EGCG concentration of ∼500 µM was used for corneal treatment. Because EGCG content was not analytically quantified, concentrations are reported as estimates rather than absolute molar doses.

### 2.2 Corneal Cultures

4 Goat (*Capra hircus*) and 8 sheep (*Ovis aries*) eyeballs were procured from a local abattoir within one hour of slaughter. Immediately after collection, the globes were gently wiped with sterile gauze soaked in phosphate-buffered saline (PBS) and subsequently rinsed in transport medium consisting of PBS supplemented with 1× penicillin–streptomycin and 1× amphotericin B. For transport, the eyes were placed individually in sterile sealed Ziploc® bags containing the transport medium. Bags were placed into insulated ice boxes for delivery to the lab, with care taken to prevent direct contact of the tissue with ice. All samples arrived for dissection within two hours of the initial decontamination. Upon arrival, all subsequent handling was performed under aseptic conditions in a biosafety cabinet (BSL-2). Each globe was removed from its transport bag and dissected individually to isolate corneoscleral rings using sterile surgical instruments. The dissected rings were immediately transferred into individual Petri dishes containing sterile PBS, and washed two more times with PBS supplemented with 1× penicillin–streptomycin and 1× amphotericin B.

Corneoscleral rings were positioned epithelial side up on agarose–gelatin solid supports placed in 6-well plates. This orientation ensured that the endothelial layer remained in contact with the supporting dome while the epithelial surface was exposed to air, mimicking an air– liquid interface (ALI) model. Each well was filled with culture medium up to the limbal boundary, with additional medium applied dropwise over the epithelial surface to maintain hydration. Cultures were maintained at 37 °C in a humidified 5% CO_2_ incubator. The medium was refreshed daily, and epithelial hydration was supplemented at each media change.

### 2.3 Assessment of *Ex Vivo* Tissue Viability

To establish goat and sheep corneas as feasible *ex vivo* models for enzymatic challenge, we performed a preliminary tissue viability assessment under our air–liquid interface (ALI) culture conditions. Corneal tissue viability was assessed using macroscopic and histological endpoints, including transparency and general gross tissue appearance. The objective was to document short-term tissue stability and integrity prior to enzymatic challenge, rather than measuring metabolic viability. We monitored transparency and contamination daily and obtained paired histology at baseline (Day 0) and after culture (Day 7) to quantify structural changes. Macroscopic observations of media color and transparency were taken every day, and the wells were observed under an inverted microscope for contamination.

**Figure 1.**
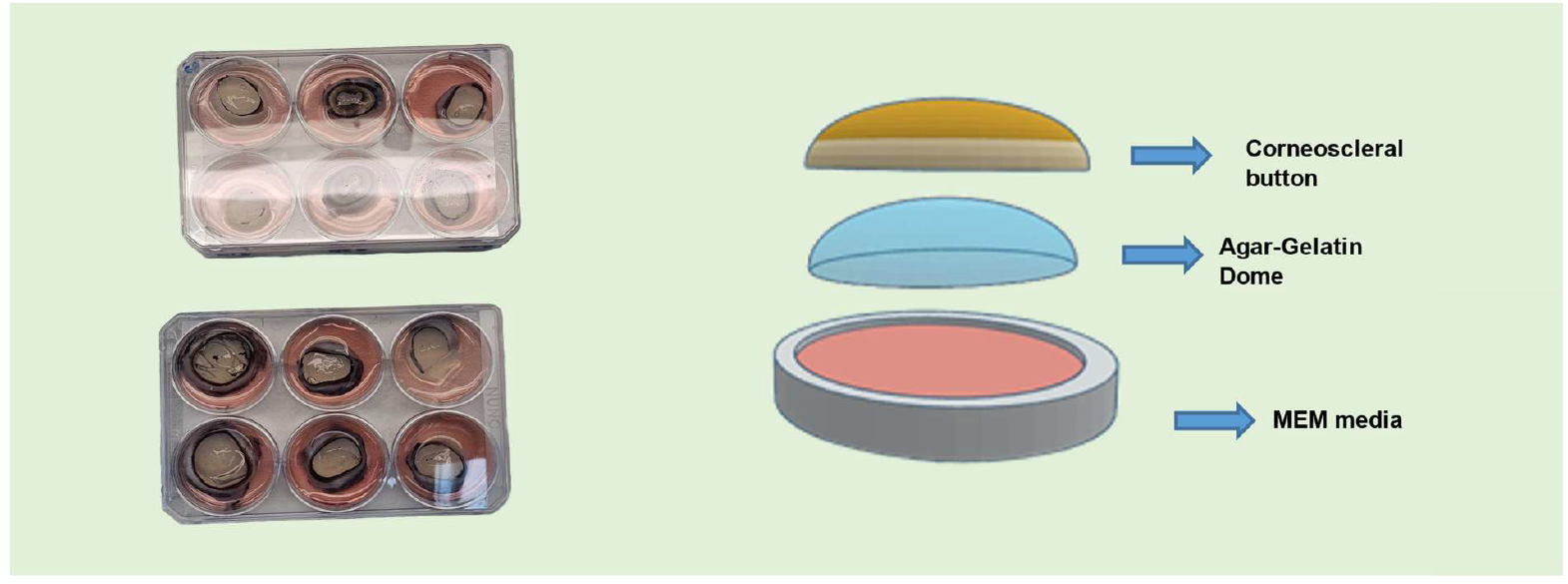
A) Figures showing the Corneoscleral Buttons (CSRs) after dissection (on the left) and in the culture medium inside a 6-well plate (on the right). B) Diagram showing the air-liquid Interface (ALI) for *ex vivo* corneal culturing. It consists of the CSRs being spread on concave agarose-gelatin domes, which are placed directly inside the culture medium, such that the epithelium (upper side) of CSRs is exposed to the air, whereas the endothelium (lower side) receives the nutrients directly, mimicking the natural eye makeup

**Figure 2.**
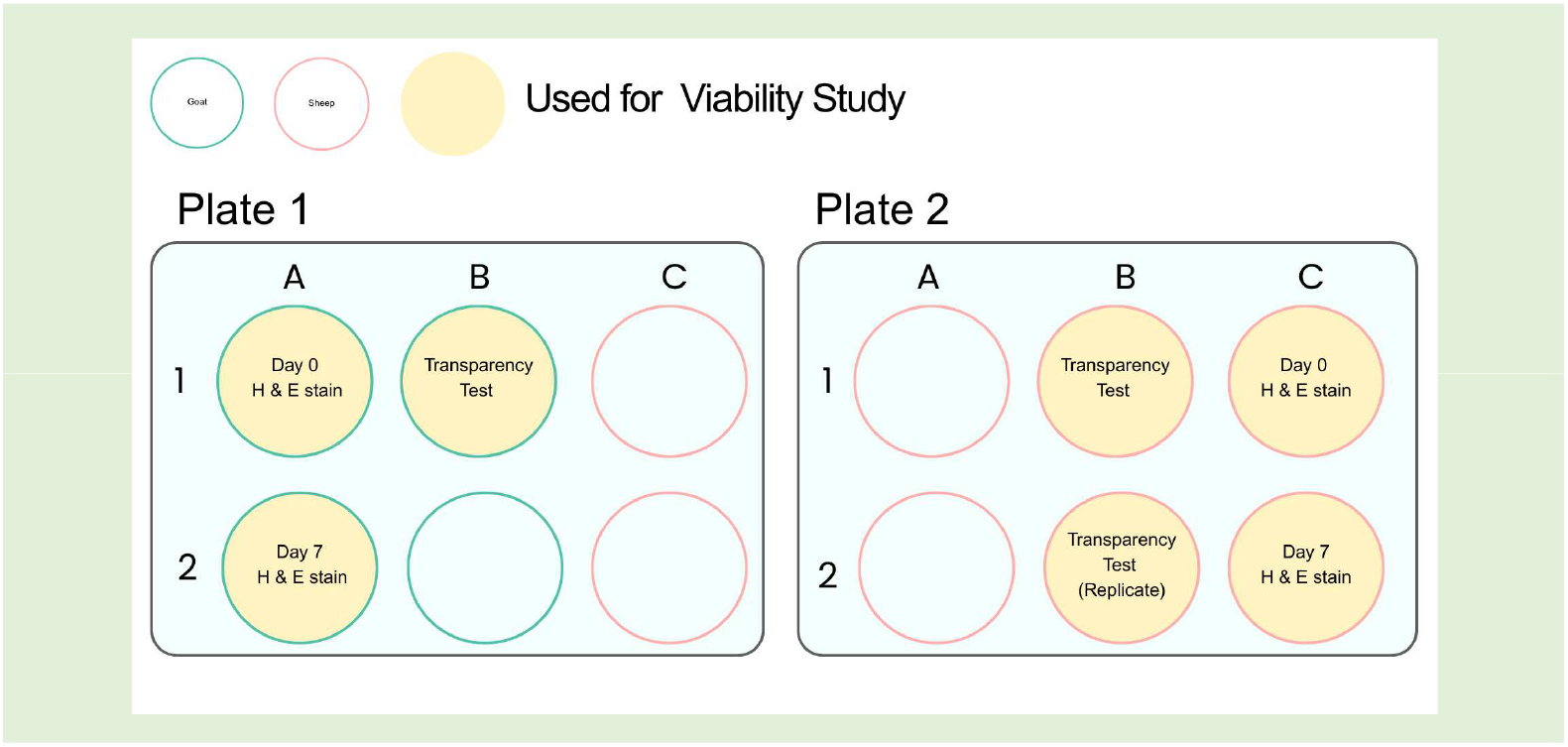
Schematic of well-plate allocation for preliminary viability study. Plate 1 was used for goat eyes (A1, A2, B1) and Plate 2 for sheep eyes (B1, B2, C1, C2). Wells were assigned to Day 0 H&E, Day 7 H&E, and transparency monitoring, as indicated.

#### 2.3.1 Transparency Grading (macroscopic)

Corneoscleral rings (CSRs) were placed in Petri dishes containing 1 mL PBS and positioned over a printed “AA” (11-point Arial font) target (Marlo *et al*. 2017). Two independent observers graded optical clarity using a 4-point rubric: 0 = fully transparent (font sharp), 1 = slight haze (font readable), 2 = moderate haze (font partially visible), 3 = opaque (font not visible). Grading was performed once every 2 days at fixed lighting and camera distance. For analysis, we averaged observers’ scores per sample.

### 2.4 EGCG Treatment Study Design

#### 2.4.1 Optimal Collagenase dosage and concentration

To determine the optimal dosage and concentration of collagenase type I used, a pilot study was run on day 5. One Sheep CSR was selected at random (A2). The CSR was taken out along with the agarose-gelatin dome and de-epithelialized by rubbing its surface gently with 20% (v/v) alcohol for 15 seconds. Then, with sterile forceps, the cornea was divided into 4 quadrants, put into 12-well plates inside their randomly assigned concentrations of collagenase type I, and then placed inside a non-CO_2_ microbiology incubator at 31 °C. Collagenase type I was prepared as % (w/v) solutions in phosphate-buffered saline (PBS) supplemented with calcium ions, which are required for collagenase activity. Concentrations tested in the pilot study were 0.0%, 0.1%, 0.3%, and 0.5% (w/v), corresponding to 0, 1, 3, and 5 mg/mL, respectively. Gross microscopic observations and light microscopy of CSR quadrants inside the wells were taken every 30 minutes, and the approximate time for dissolution, along with visible surface morphology features, were noted. Concentration of 0.3% yielded consistent, time-bound digestion (complete breakdown within the planned window, 4– 5 h) and was selected as the challenge concentration for subsequent EGCG experiments.

#### 2.4.2 EGCG Experimental Study

To find the protective effects of EGCG extract on corneal tissues, an experimental study was conducted on the last day (Day 7) of the culturing protocol. CSR C1 was used for sheep, and B2 for goats. After de-epithelialization, the CSRs were quartered, and the four quadrants were randomized to pretreatments inside 4 wells of a 12-well plate.

Two quadrants were incubated in 500 microliters of ∼500 µM EGCG extract solution for 1.5 h at 30–31 °C. Plates were wrapped in aluminum foil with the incubator lights switched off to minimize light-induced degradation of EGCG. Whereas the remaining two quadrants were incubated in 500 microliters of PBS for 1.5 h at 30–31 °C (negative control). Following pretreatment, quadrants were thoroughly rinsed in PBS (x3) and moved to their post-treatments for 4.5 hours, as determined in the pilot. In post-treatments, each corneal quadrant was incubated in 500 µL of collagenase solution per well or equivalent volumes of EGCG or PBS. Details of treatment allocations are summarized in Table 2.

**Table 1.** This table summarizes the potential therapeutic effects of EGCG in the context of Keratoconus treatment.

| Effect | Mechanism | Evidence Source | Relevance to KC | Limitations |
| --- | --- | --- | --- | --- |
| **Structural Cross-linking** | EGCG cross-links collagen fibrils, increasing stiffness | Elder *et al.*, 2017; Chu *et al.*, 2016; Sapula *et al.*, 2023 | Could reinforce stromal biomechanics, similar to CXL | Evidence mostly in bone, dentin, xenografts, hydrogels, not cornea |
| **Resistance to Enzymatic Digestion** | Cross-linked collagen resists collagenase/MMP breakdown | Chu *et al.*, 2016 | Protects stromal fibrils against degradation | *In vitro*, non-corneal |
| **Anti-inflammatory** | Downregulates IL-1β, TNF-α, and MMP-9 pathways | Cavet *et al.*, 2011; Urolita *et al.*, 2024 | Addresses KC-associated inflammatory upregulation | Mostly keratitis/epithelial models |
| **Antioxidant** | Scavenges ROS, protects epithelial/stromal cells | Capasso *et al.*, 2025; Ng *et al.*, 2023 | Counteracts oxidative stress, a major KC factor | Preclinical, not KC-specific |
| **Wound Healing** | Promotes re-epithelialization, reduces scarring | Urolita *et al.*, 2024 | Supports epithelial integrity, reduces KC complications | Acute wound models may not recapitulate chronic KC pathobiology |
| **Collagenase Inhibition** | Direct inhibition of collagenase and MMP activity | Chu *et al.*, 2016; Urolita *et al.*, 2024 | Protects stromal collagen against enzymatic thinning | Needs direct stromal validation |

**Table 2.** This table summarizes the pre-treatment and post-treatment allocations for each CSR Quadrant.

|  |  |  |
| --- | --- | --- |
| 1. | EGCG → COL: Pretreated with EGCG, then exposed to collagenase. | Tests whether EGCG pretreatment confers resistance to enzymatic digestion. |
| 2. | PBS → COL: Pretreated with PBS, then exposed to collagenase. | Serves as the collagenase control; used to compare with EGCG → COL to assess the protective effect of EGCG pretreatment. |
| 3. | EGCG → EGCG: Pretreated with EGCG, then maintained in EGCG | Tests whether continuous EGCG exposure protects stromal integrity, compared with PBS controls. |
| 4. | PBS → PBS: Pretreated with PBS, then maintained in PBS | Serves as the negative control; used to assess baseline culture effects without EGCG or collagenase. |

**Figure 3.**
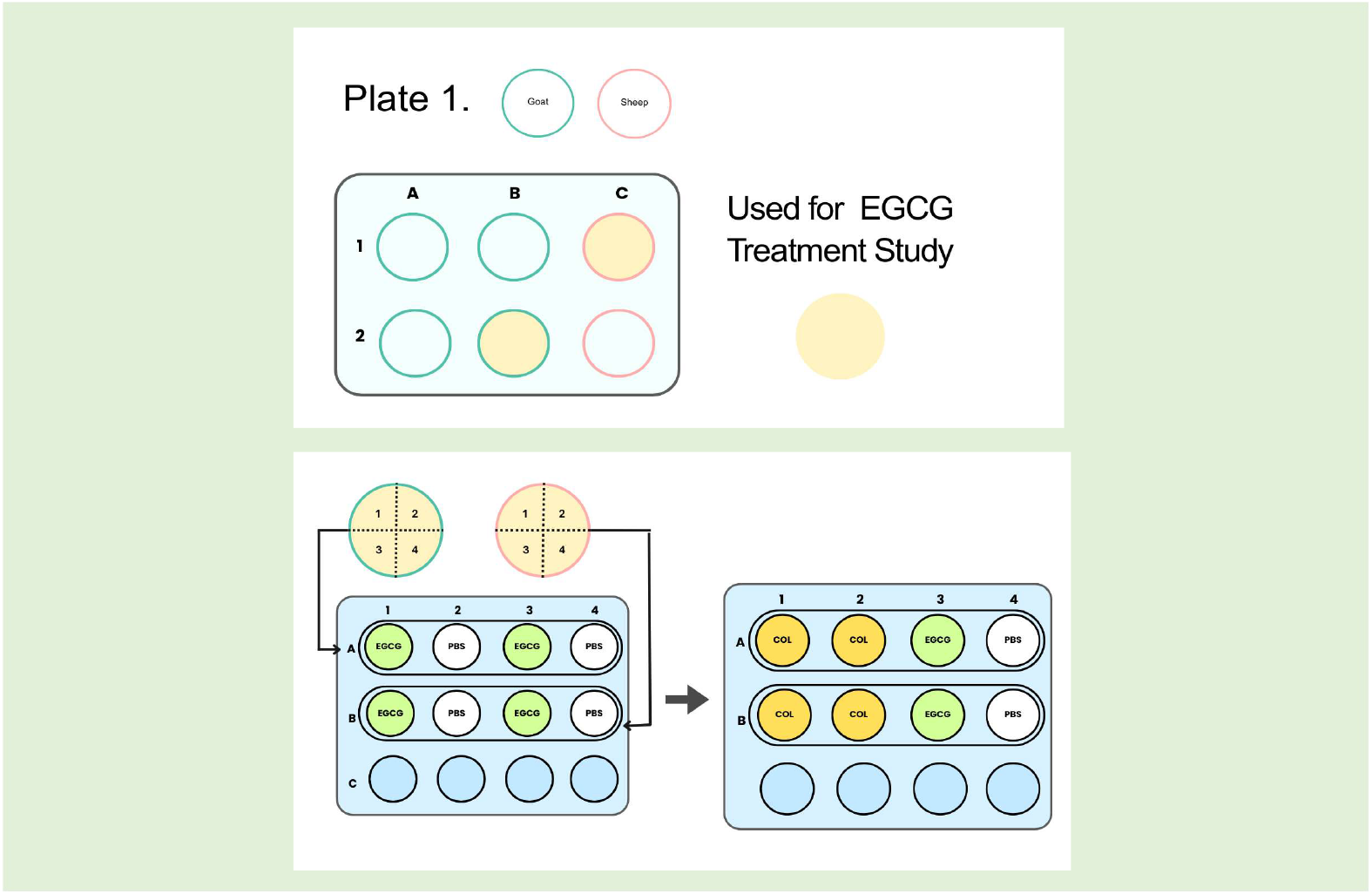
Schematic of well-plate allocation for EGCG treatment study. Each CSR was divided into four quadrants, randomized into four treatment groups: EGCG → COL: Pretreated with EGCG, then exposed to collagenase. PBS → COL: Pretreated with PBS, then exposed to collagenase. EGCG → EGCG: Pretreated with EGCG, then maintained in EGCG. PBS → PBS: Pretreated with PBS, then maintained in PBS.

All incubations used sufficient volume to fully immerse the tissue and prevent desiccation. Exposure times matched those determined in the pilot (collagenase) or pretreatment (EGCG/PBS). After 5 hours of post-treatment, the quadrants were removed, fixed in 10% neutral buffered formalin, and sent for H&E Staining. Gross Microscopic Images were taken of all samples.

#### 2.4.3 Eosin-positive area fraction analysis using H&E color deconvolution

To assess stromal preservation following enzymatic challenge, eosin-positive stromal area fraction analysis was performed on hematoxylin and eosin (H&E)–stained corneal sections using Fiji/ImageJ (ImageJ version 1.54p; National Institutes of Health, Bethesda, MD, USA), following established image-based histological quantification approaches (Walker *et al*., 2018; Naka *et al*., 2020). Eosin preferentially stains protein-rich components of the stromal extracellular matrix, including but not limited to collagen, and loss of eosin staining has been widely used as an indicator of tissue degradation in histological injury and digestion models (Walker *et al*., 2018; Naka *et al*., 2020. In H&E-stained corneal sections, intact stromal lamellae therefore appear strongly eosinophilic, whereas enzymatic degradation of stromal matrix proteins results in reduced eosin staining. Accordingly, this approach provides a semi-quantitative surrogate measure of stromal matrix integrity based on retained eosin staining within a defined tissue area, rather than a collagen-specific assessment. The eosin positive fraction assay (EMA) was run through Image Processing in Fiji to assess stromal preservation, as described by Walker *et al*. (2018). Quantitative eosin-positive area fraction analysis was performed exclusively on 40× magnification brightfield images, whereas 10× images were used for representative visualization of histological features.

**Figure 4.**
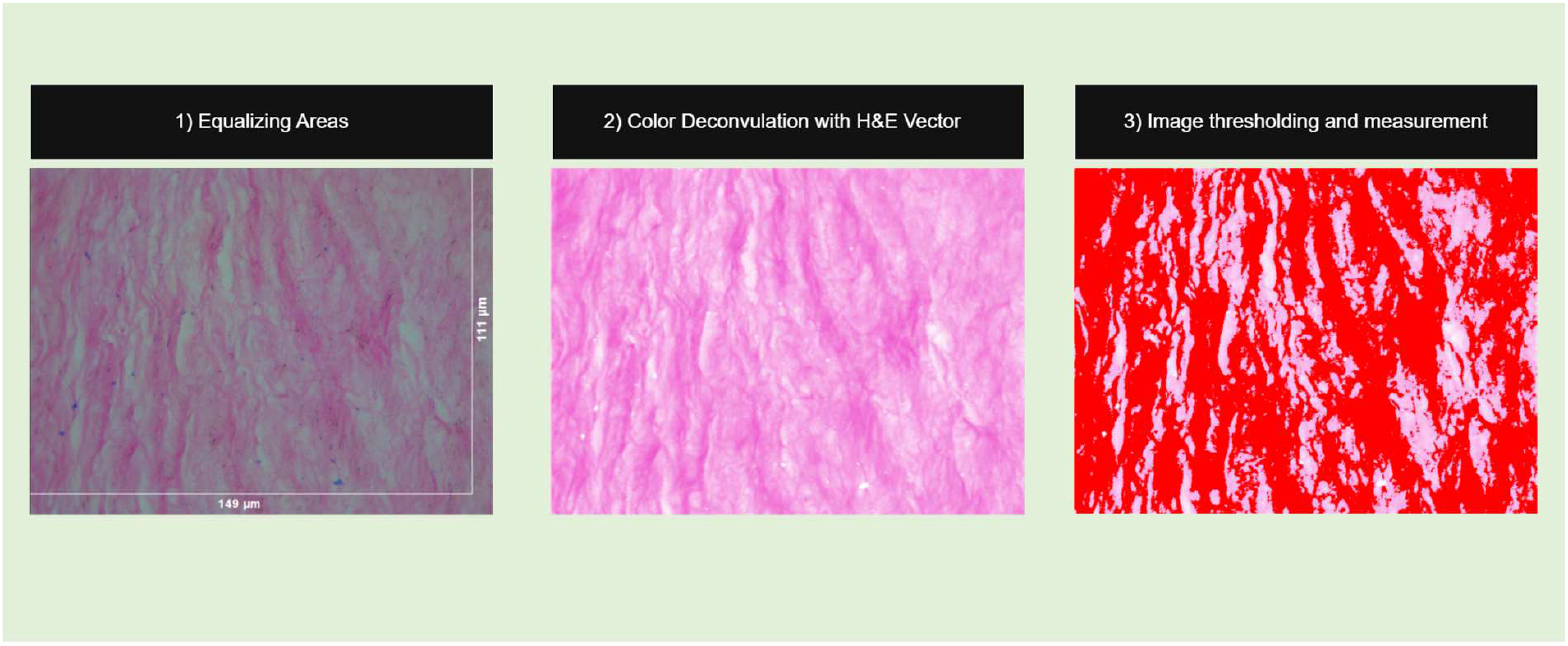
Figure demonstrating the steps used for determining the Eosin-positive area fraction for the corneal samples. For each experimental condition, three representative 40× brightfield images were acquired from the anterior stromal region of the same corneal sample, immediately beneath the epithelium. Images were imported into Fiji/ImageJ and calibrated using the microscope scale bar. A fixed region of interest (ROI) of equal area (149 × 111 µm) was applied across all images within each condition to ensure consistency. Color deconvolution using the standard H&E vector was performed to isolate the eosin channel, followed by local contrast enhancement. The eosin channel was then thresholded using the intermodal (inter-modes) method to distinguish eosin-preserved stromal regions from degraded or unstained areas. The eosin-positive area within the ROI was measured for each image, and values from three fields were averaged. Results were expressed as the percentage of eosin-positive stromal area relative to the total ROI area. In this context, higher eosin-positive area fractions reflect greater retention of protein-rich stromal extracellular matrix and overall stromal architectural integrity following enzymatic challenge.

## 3.0 Results

### 3.1 Ex Vivo Tissue Viability

The dissected CSRs were placed in 6-well plates for 7 days, to assess short-term tissue (tissue viability) prior to enzymatic challenge. This was done by macroscopic observations of the culture medium, measuring the transparency grade of the corneas as well as conducting histological evaluation on prepared slides of the corneal tissue (Day 0 and Day 7). Gradual loss of transparency was observed with most tissues appearing hazy and visibly opaque by Day 7. Beyond that point, the CSRs were not cultured. The 2 CSRs who developed premature whitening due to contamination were excluded. To minimize contamination in samples that were retained, the concentration of penicillin-streptomycin and amphotericin B was increased from 1x to 2x inside the culture medium on Day 3. Uncontaminated corneas were successfully carried through to Day 7, beyond which they were not cultured.

#### 3.1.1 Transparency Grading

Transparency grades of the CSRs were obtained by two independent observers and averaged per cornea at each interval. The important thing to note is that, on day 5 observations, culture medium spread over the corneal surface, exaggerating the opacity: multiple PBS washes revealed the underlying transparency level. The same was done for day 7; however, even after multiple washes, the print was not visible.

**Figure 5.**
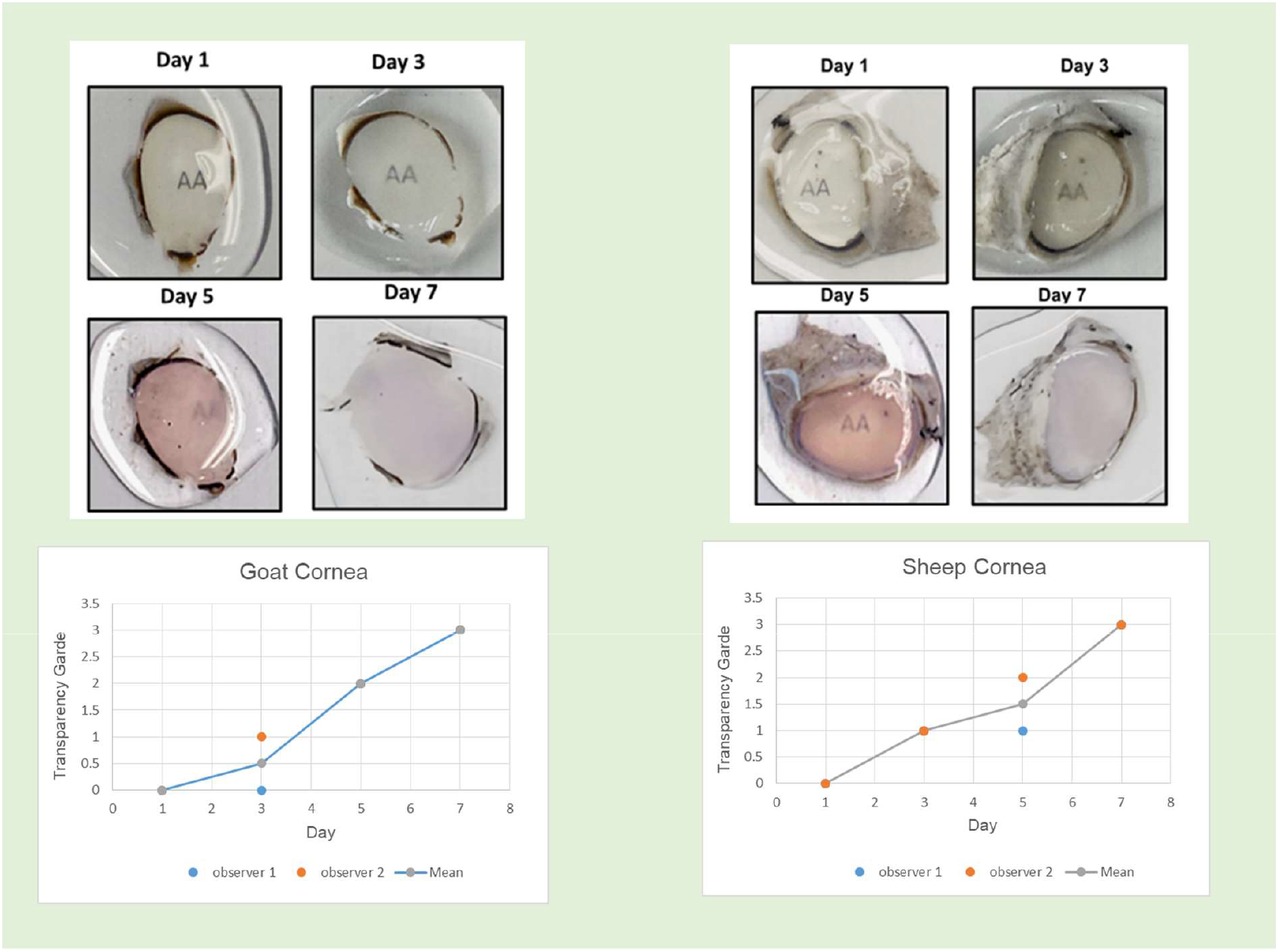
On day 1, both Goat and Sheep corneas were fully transparent, with the font being fully legible. On day 3, a very mild haze was evident with goat corneas averaging grade 0.5 and sheep slightly hazier at 1. By day 5, transparency had deteriorated further: goat corneas reached grade 2 (moderate haze, font only slightly visible), whereas sheep corneas averaged 1.5 (between slight and moderate haze). By day 7, both species had become opaque, with a mean grade of 3, and the underlying print was no longer visible.

### 3.2: EGCG treatment Study

**Figure 6.**
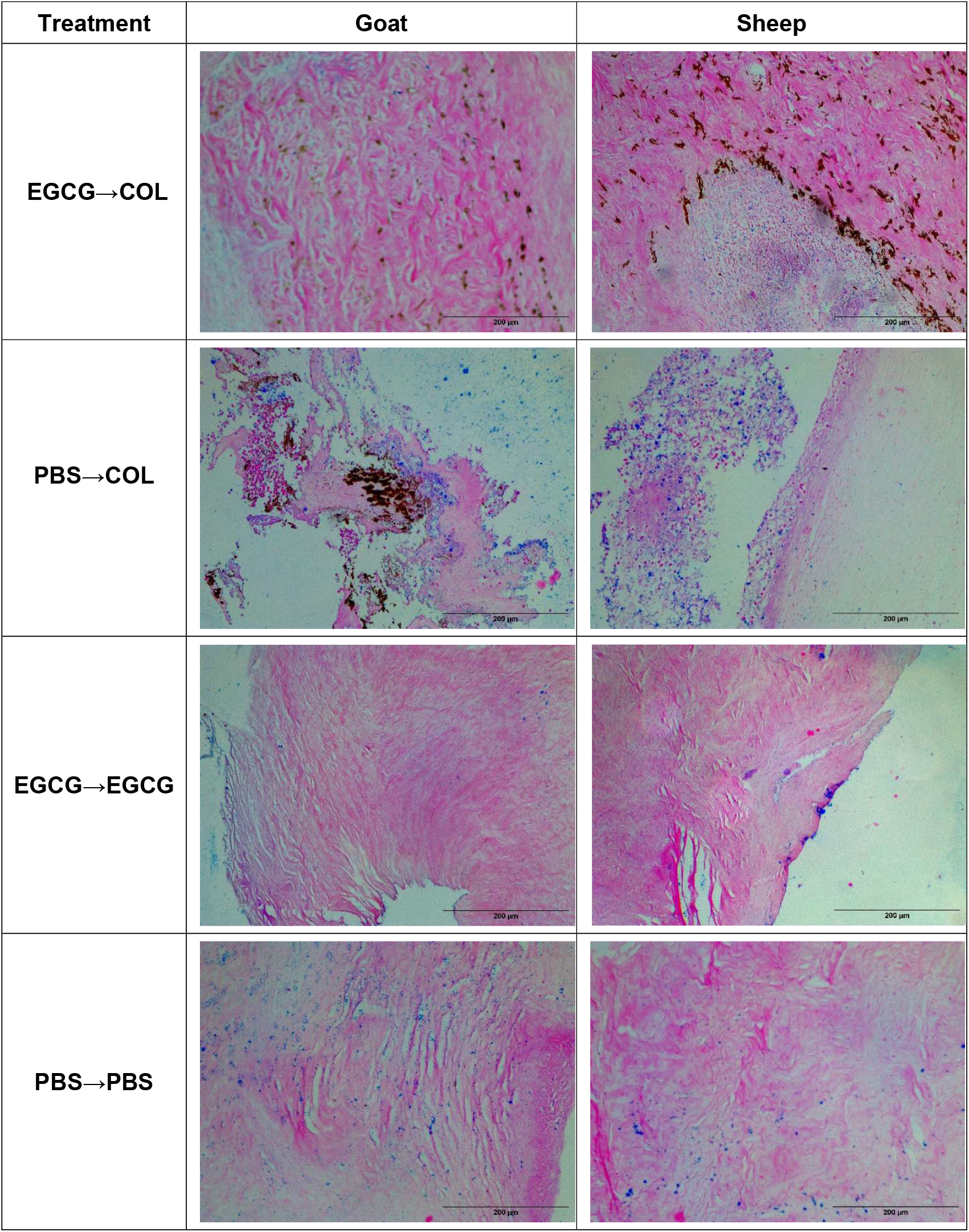
Representative histological samples for each treatment group. The samples were stained with H&E and captured at 10x magnification for gross evaluation. Scale bar = 200 µm.

#### 3.2.2 Qualitative histological observations and quantitative eosin-positive stromal area fraction

**Table 3.**
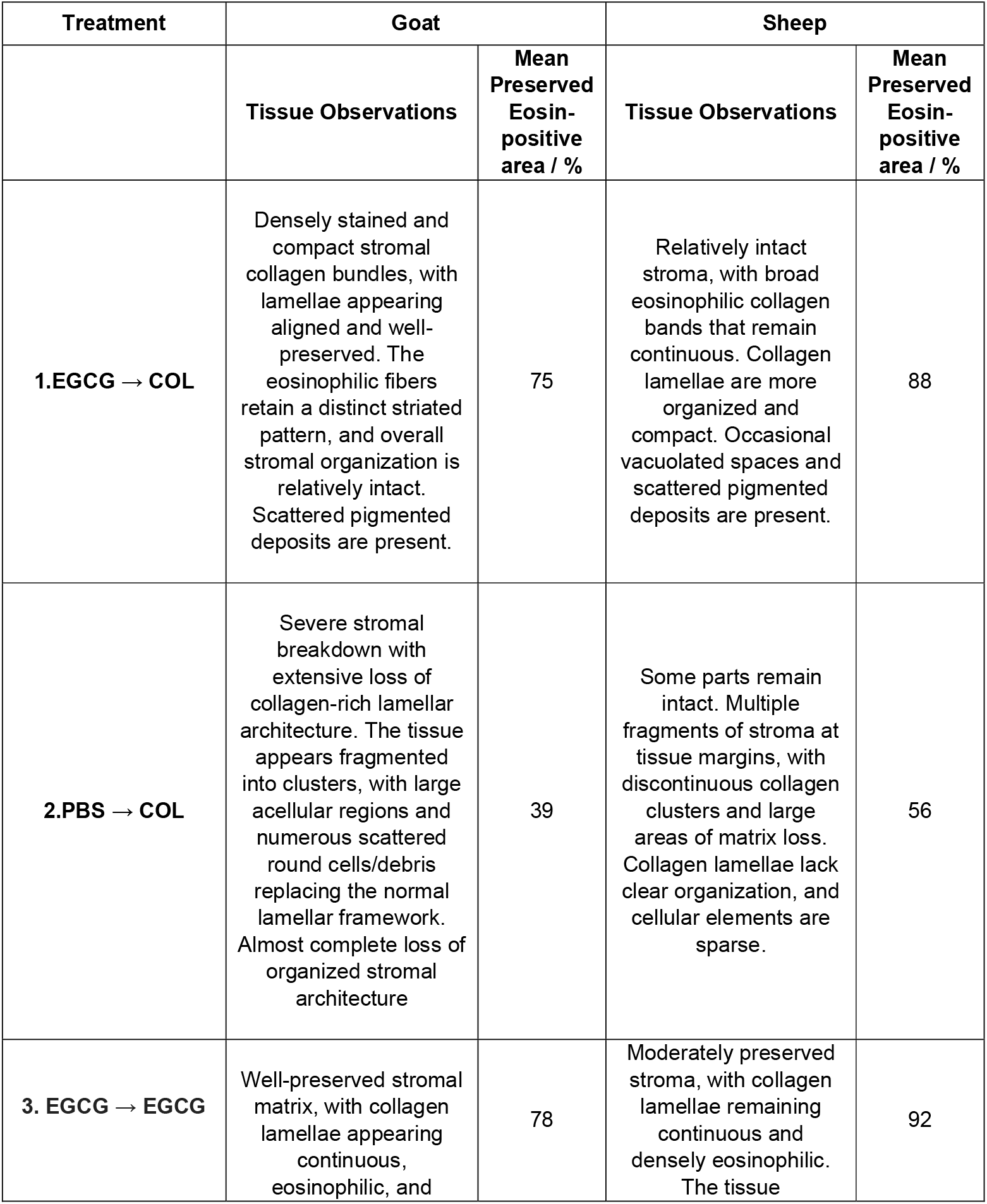
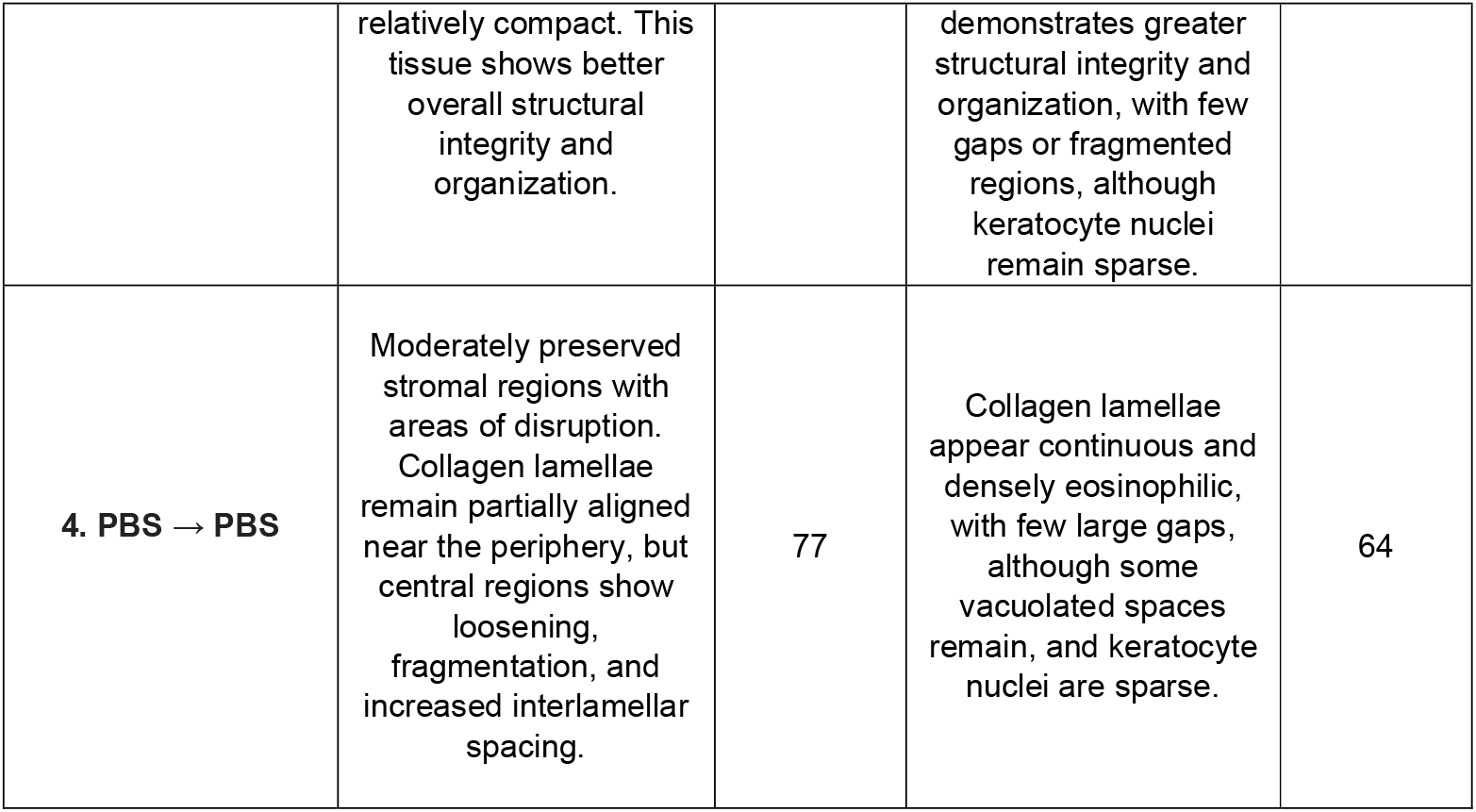
This table represents the blind observations of the H&E-stained slides for representative goat and sheep quadrants following EGCG and collagenase treatments. Observations of cellular elements, lamellar appearance, interlamellar spaces, and overall stromal organization were noted. Adjacent columns represent the preserved stromal fraction, quantified by eosin staining. Reported eosin-positive area fractions (%) represent measurements from representative quadrants derived from a single cornea per species and are intended as semi-quantitative, descriptive indicators rather than statistically powered estimates. A high eosin-stained stromal area represents greater stromal matrix preservation.

## 4.0 Discussion

### 4.1 EGCG as a therapeutic for KC

This preliminary study demonstrates that EGCG extract pretreatment was associated with stromal preservation in goat and sheep corneal stroma from collagenase-induced degradation. Furthermore, the EGCG extract pre-treated and post-treated corneas had the greatest relative preservation of stromal architecture, even more than PBS controls, as supported by the histological observations and quantification of eosin-stained stromal area.

For Goat CSR, PBS→COL treatment completely dissolved the CSR quadrant as was observed macroscopically, consistent with the lowest preserved eosin-stained stromal area (39%). Under the microscope, a complete loss of stromal architecture was observed, denoting the goat cornea’s high susceptibility towards digestion by collagenase. However enzymatic challenge, with EGCG extract pretreatment (EGCG → COL), yielded notably converse results. The stroma was in one piece, with compact collagen bundles and aligned lamellae (75% preserved area). Although the stromal ultrastructure had a striated pattern, which could show part or differential digestion. Notably, all the CSRs were de-epithelialized before, and a complete loss of stroma organization vs preserved stroma in the above condition demonstrated plausible penetration of the respective treatments. More so, in PBS→PBS goat CSR, eosin staining showed 77% preserved stromal area, but large interlamellar gaps were evident, likely from stromal swelling, consistent with better penetration in goat tissue.

For sheep CSR, PBS→COL caused less extensive digestion (56% preserved area), with regions of intact stroma remaining. This partial preservation likely reflects the greater size and thickness of sheep corneas, which limit enzyme penetration. Still, EGCG→COL maintained the highest stromal preservation (88% area), with less fragmentation. In PBS→PBS sheep CSR, preservation was lower (64%) compared to goat (77%), again reflecting differences in tissue swelling and thickness. It is important to note that some CSRs displayed sharp stromal breaks from mechanical handling; these were readily distinguishable from collagenase-mediated digestion, which produced more diffuse and uniform stromal degradation.

These findings demonstrate that EGCG confers resistance to collagenase-mediated stromal degradation. Mechanistically, EGCG’s protective effect is likely two-fold: it may directly fortify the collagen matrix via chemical cross-linking, and it may inhibit collagen-degrading enzymes in the corneal tissue. Both mechanisms would result in reduced collagen digestion and better structural integrity, aligning with the observed preservation of the corneal stroma in this *ex vivo* model.

EGCG’s ability to cross-link and strengthen collagen has been reported in tissue engineering contexts. For example, Chu *et al*. (2016) found that collagen membranes treated with EGCG formed stable cross-links (Schiff bases) without altering the collagen’s ultrastructure, effectively increasing resistance to collagenase degradation. Similarly, a recent review by Sapula *et al*. (2023) concluded that EGCG cross-linking of biopolymer hydrogels confers greater resistance to enzymatic breakdown and improves mechanical stiffness, while maintaining high biocompatibility and low cytotoxicity. These properties are consistent with our results: EGCG-treated corneas resisted enzymatic digestion, and support the concept that EGCG may act as a stromal stabilizer in corneal tissue, potentially through collagen interactions and/or enzyme inhibition. (Capasso *et al*. 2025).

Notably, EGCG also exhibits anti-inflammatory effects in ocular tissues, which could further contribute to corneal preservation. In keratitis models, Urolita *et al*. (2024) reported that EGCG suppresses IL-1β–induced NF-κB activation and downstream uPA/MMP-9 expression in corneal fibroblasts, thereby reducing the enzymatic collagen degradation that inflammatory cells can cause. Cavet *et al*. (2011) similarly showed that EGCG acts as a potent anti-inflammatory and anti-oxidant agent in human corneal epithelial cells, dose-dependently inhibiting IL-1β–stimulated release of inflammatory cytokines without any cytotoxic effect. These literature findings suggest that EGCG could mitigate the inflammatory cascades (e.g. MMP-9 upregulation) associated with corneal injury or Keratoconus. This dual action – cross-linking collagen and dampening inflammation – is particularly valuable, as chronic Keratoconus progression and corneal ulceration involve both biomechanical weakening and inflammatory matrix degradation. The protective outcome aligns well with the above studies and indicates that EGCG addresses both the structural and inflammatory aspects of corneal degeneration.

Given these promising results, it is important to compare EGCG’s potential therapeutic role to the current standard of care for Keratoconus - riboflavin/UVA cross-linking (CXL). While CXL effectively halts keratoconus progression, it carries notable limitations, including dependence on corneal thickness (>400 μm) and risks of pain, haze, infection, and endothelial damage. In contrast, EGCG would not be constrained by corneal thickness or require UV irradiation, making it potentially safer for thin or advanced keratoconic corneas. Furthermore, EGCG’s anti-inflammatory activity could be especially valuable in inflamed or high-risk corneas, offering dual protection by reinforcing stromal collagen and attenuating biochemical drivers of degeneration.

From a safety perspective, cytotoxicity studies have defined a therapeutic window for EGCG in ocular tissues. Corneal fibroblasts and epithelial cells remain viable at concentrations below 30 µM, whereas apoptosis and viability loss emerge at doses ≥100 µM, highlighting the importance of careful dosing (Sugioka *et al*., 2019; Cavet *et al*., 2011). Animal studies have further demonstrated that topical formulations up to 0.1% EGCG do not induce significant epithelial toxicity, supporting the feasibility of ocular application (Huang *et al*., 2018). However, it is important to recognize that the concentration used in this study (∼500 µM) is above these reported thresholds, reinforcing that while *ex vivo* experiments establish proof-of-concept, future clinical translation will require optimization of delivery systems to achieve efficacy within physiologically tolerable ranges.

Penetration across the corneal epithelium is another critical consideration. The epithelium forms a significant barrier to stromal delivery of hydrophilic compounds like EGCG, which is the same limitation that necessitates epithelial removal (“epi-off”) in conventional riboflavin/UVA CXL to allow adequate riboflavin penetration. Formulation advances provide promising alternatives, as nanoparticle and gelatin-based carriers have been shown to prolong corneal residence time, enhance stromal penetration, and broaden ocular distribution while avoiding epithelial toxicity (Huang *et al*., 2018). Mechanistic data also indicate that EGCG suppresses IL-1β–driven uPA/MMP-1 cascades in corneal fibroblasts, suggesting that even partial stromal penetration may suffice to modulate pathogenic enzymatic activity (Sugioka *et al*., 2019). Together, these findings support the feasibility of EGCG as a topical or minimally invasive therapy, provided that dosing and delivery are carefully tailored to balance efficacy with safety.

### 4.2 *Ex Vivo* Models as a mode for Ocular disease modelling and drug testing

A key objective of this study was also to investigate goat and sheep corneas as practical *ex vivo* models for corneal ectasia and its treatments. The culture methodology adopted from several *ex vivo* corneal studies (Madhu et al., 2018; Marlo et al., 2017) successfully maintained goat and sheep corneoscleral rings in an air–liquid interface for 7 days. Early in the culture period, some microbial contamination was observed; this was controlled by increasing the concentrations of penicillin–streptomycin and amphotericin B in the medium. While effective, high levels of antimicrobials may also compromise tissue health by altering nutrient availability or exerting cytotoxic effects on resident stromal cells. Daily monitoring showed progressive haze development, and by Day 7 the corneas demonstrated complete whitening (loss of optical transparency), indicating that although stromal architecture could still be assessed histologically, functional optical clarity was no longer maintained.

The ability to sustain corneas for a week is sufficient for simulating enzymatic stromal degradation and evaluating short-to-intermediate term therapies such as cross-linkers or drug delivery systems. The air–liquid interface (ALI) model we employed (with the endothelium bathed in media and epithelium exposed to air) closely mimics physiological conditions and has been used successfully in other species’ *ex vivo* corneas. By preserving native cell–matrix interactions and avoiding supra-physiological swelling, the ALI culture helps maintain normal stromal ultrastructure over time. This approach thus provides a robust platform to study short-term corneal ectatic changes and interventions under controlled laboratory settings, without the ethical and logistical complexities of *in vivo* animal experiments.

When considered specifically for *ex vivo* corneal culturing, goat and sheep corneas offer several advantages over other models. Small animals such as mice and rats are useful for genetic studies, but their corneas are too thin (∼100 μm) and small for practical biomechanical or surgical testing (Wei *et al*., 2023). Rabbits are commonly used *in vivo* corneal research, but their use demands specialized animal facilities and raises ethical concerns, especially since many experiments require euthanasia. In contrast, goat and sheep eyes can be sourced ethically and cost-effectively from abattoirs as discarded tissue, eliminating the need for purpose-bred animals (Madhu *et al*., 2018). Anatomically, goat corneas (∼600–650 μm thick) and sheep corneas (∼700 μm thick, with globes ∼30% larger than human eyes) are both closer in size and handling characteristics to human corneas. Importantly, sheep corneas possess a Bowman’s layer, which may make their wound-healing and biomechanical responses more comparable to humans (Shankar *et al*., 2024). Proteomic analyses also show ∼90% similarity between sheep and human corneal proteins, suggesting biochemical comparability at the molecular level. Although porcine and bovine corneas are also widely used in cross-linking and surgical training studies, cultural and religious restrictions in South Asia and the Middle East limit their accessibility. Goat and sheep corneas therefore occupy an intermediate niche: they are large enough for surgical and biomechanical testing, logistically simple to obtain, and culturally acceptable, yet underutilized in corneal research.

Our successful use of goat and sheep corneas *ex vivo* demonstrates their potential as reliable models for short-term culture and experimentation. While issues such as antimicrobial toxicity, initial contamination control, and medium selection remain important limitations, these do not diminish the value of such models in advancing corneal research. In contexts such as Pakistan, where access to human donor tissue or porcine models is limited, leveraging abattoir-sourced goat and sheep eyes could greatly accelerate translational corneal studies, providing an ethical and cost-effective platform for drug screening, cross-linker testing, and tissue engineering applications in alignment with the 3Rs (Replacement, Reduction, Refinement) of animal research ethics.

### 4.3 Limitations

As a preliminary *ex vivo* investigation, this study has several inherent limitations that should be considered. First, the sample size was very small, and the findings should be regarded as preliminary until validated in larger cohorts that capture inter-animal variability. Second, the *ex vivo* nature of the experiments, while useful for controlled mechanistic testing, does not recapitulate the systemic, immune, and wound-healing responses that would occur *in vivo*. Third, tissue integrity was assessed primarily through histology and eosin quantification without biomechanical testing, so the degree of stromal stiffening conferred by EGCG remains unknown. Additionally, because EGCG is a known collagenase and matrix metalloproteinase inhibitor, and because tissue-associated EGCG may persist despite PBS washing, this study cannot definitively distinguish between stromal protection mediated by direct enzymatic inhibition versus permanent collagen cross-linking. Finally, the concentration of EGCG in the extract used (500 µM) is above the safe therapeutic window reported for corneal cells (≥100 µM). While this high dose effectively demonstrated proof-of-concept enzymatic resistance, future studies must optimize dosing and employ controlled-release strategies to achieve efficacy within physiologically tolerable ranges.

Culture-related limitations also warrant consideration. Minimum Essential Medium (MEM) was employed due to reagent availability constraints, and its lower baseline nutrient content was partially mitigated through supplementation with fetal bovine serum and epidermal growth factor (EGF) to support short-term keratocyte cell viability by delaying stress-induced apoptosis during ex vivo culture. However, the use of MEM rather than Dulbecco’s Modified Eagle Medium (DMEM) likely limited tissue viability to 7 days, as DMEM provides higher concentrations of amino acids, vitamins, and glucose that are better suited to sustaining long-term corneal culture. Early microbial contamination required supplementation with higher concentrations of penicillin–streptomycin and amphotericin B, which, while effective, may have reduced nutrient availability or imposed cytotoxic stress on stromal cells. A probable contributing factor was the absence of povidone-iodine (betadine) sterilization. This may explain why our models were limited to 7 days of stability, whereas other studies using DMEM and betadine sterilization have reported goat corneal viability up to 15 days in culture (Madhu *et al*., 2018). By Day 7, all corneas exhibited complete whitening and reduced keratocyte density, meaning that stromal ultrastructure could be studied histologically, but functional optical clarity was lost. These limitations highlight that while goat and sheep corneas provide a practical and underutilized platform for *ex vivo* research, improvements in medium formulation, contamination control, and tissue handling are necessary to maximize their reliability for longer-term studies.

## 5.0 Conclusion and Future Directions

In conclusion, we tested the effects of epigallocatechin gallate (through an EGCG-rich extract) in a collagenase-induced *ex vivo* model of corneal stromal degradation using caprine and ovine corneas. EGCG extract pretreatment was associated with notably greater preservation of stromal architecture and eosin-stained stromal architecture compared with collagenase-only and PBS controls. In parallel, this study establishes goat and sheep corneas as feasible short-term *ex vivo* models for investigating enzymatic mimicking of corneal ectasia and stromal protective agents. Together, these findings demonstrate that EGCG confers resistance to collagenase-mediated stromal degradation, and has the potential of being adjunct to conventional corneal collagen cross-linking, while supporting the use of caprine and ovine corneas as practical *ex vivo* models for translational corneal research

Moving forward, several research directions are required to validate and extend the findings of this preliminary study. First, *in vivo* testing of EGCG should be prioritized in models of keratoconus or enzymatic corneal thinning (e.g., collagenase-induced ectasia in rabbits or sheep). Such studies would allow evaluation of EGCG in a physiological environment that includes keratocytes, epithelial and endothelial layers, tear film interactions, and host immune responses. This will be essential to assess potential side effects, such as epithelial toxicity or inflammatory modulation, which cannot be captured in an *ex vivo* setting.

Secondly, the cross-linking capacity of EGCG should be validated in corneal tissue using biomechanical testing and thermal stability assays (e.g., collagen shrinkage or denaturation temperature), to distinguish permanent stromal stabilization from transient enzymatic inhibition as the primary mechanism of protection. Quantitative approaches such as stress–strain testing or Brillouin microscopy could provide direct measures of stromal stiffening, complementing histological preservation. Moreover, dose–response studies are needed to determine the concentration thresholds at which EGCG achieves effective stromal stabilization or cross-linking, if present, without inducing cytotoxicity. Further exploration of delivery strategies, including topical formulations, iontophoresis, or nanoparticle-based sustained release, would help define how therapeutic stromal concentrations can be achieved while remaining within a safe biological window. Comparative studies of EGCG alone versus EGCG combined with standard corneal collagen cross-linking (CXL) may also reveal synergistic effects, such as reducing CXL-induced haze or potentially enabling effective treatment at lower UVA energy.

Finally, for the *ex vivo* goat and sheep models, future studies should focus on extending tissue viability beyond one week by optimizing culture media. Longer-term *ex vivo* viability would permit evaluation of disease progression and extended dosing regimens. Just as pig and rabbit corneal models have standardized culture protocols, species-specific parameters for goat and sheep corneas must be defined and standardized, including optimal medium composition, storage conditions, and viability assessment metrics.

## Competing interests

The authors declare that there are no relevant financial, relational, or other conflicts of interest associated with this manuscript or the research conducted.

## Author Contributions

Conceptualization, M.U.A.K. and S.T.A.; design of the study, M.U.A.K., S.T.A., and S.N.; conduct of the study, M.U.A.K, S.N. and R.N.K.; writing—original draft preparation, M.U.A.K., S.T.A.; writing—review and editing, M.U.A.K., S.T.A., S.N., R.N.K, and F.F.K.; supervision, S.T.A., S.N., and F.F.K. All authors have read and agreed to the uploaded version of the manuscript.

## Acknowledgements

The authors are grateful to the Precision Medicine Lab (PML) and Rehman Medical Institute, Peshawar, for providing the resources and accommodation for this study. We also thank Bilal Rehman, Dr. Madina Shirdel, Arsalan Riaz, Dr. Maria Tasneem Khattak, Dr. Ayesha Jamil, and Mr. Zulfiqar for their dedicated support, access to experimental facilities, and valuable assistance throughout the course of this study.

## Data Availability

The datasets generated and analyzed during the current study are available from the corresponding author upon reasonable request. Due to the nature of the study, some data are presented in the form of images and histological observations, which are available as supplementary files or upon direct inquiry.

## Notes

### Competing Interest Statement

The authors have declared no competing interest.

